# Efficient and accurate sequence generation with small-scale protein language models

**DOI:** 10.1101/2023.08.04.551626

**Authors:** Yaiza Serrano, Sergi Roda, Victor Guallar, Alexis Molina

**Affiliations:** Nostrum Biodiscovery S.L., 08029, Barcelona, Spain; Barcelona Supercomputing Center (BSC), 08034 Barcelona, Spain; Institució Catalana de Recerca i Estudis Avançats (ICREA), 08010 Barcelona, Spain

**Keywords:** Protein design, Enzyme engineering, Protein language models

## Abstract

Large Language Models (LLMs) have demonstrated exceptional capabilities in understanding contextual relationships, outperforming traditional methodologies in downstream tasks such as text generation and sentence classification. This success has been mirrored in the realm of protein language models (pLMs), where proteins are encoded as text via their amino acid sequences. However, the training of pLMs, which involves tens to hundreds of millions of sequences and hundreds of millions to billions of parameters, poses a significant computational challenge.

In this study, we introduce a Small-Scale Protein Language Model (SS-pLM), a more accessible approach that requires training on merely millions of representative sequences, reducing the number of trainable parameters to 14.8M. This model significantly reduces the computational load, thereby democratizing the use of foundational models in protein studies. We demonstrate that the performance of our model, when fine-tuned to a specific set of sequences for generation, is comparable to that of larger, more computationally demanding pLM.

## Introduction

Proteins, the intricate biological molecules of life, are a marvel of complexity and diversity. Composed of 20 different amino acids, these linear polymers have the ability to fold into a vast array of 3D structures, each one enabling an intricate set of functions. Natural proteins arise under the light of evolution, sampling only a fraction of the theoretically possible sequence space. Even short proteins of 100 residues can have an immense number of potential sequences (20^100^)(1). Protein engineering stands as a pivotal field with profound implications across a multitude of sectors, including healthcare, agriculture, and environmental science. Its significance is rooted in its capacity to manipulate the fundamental building blocks of life. As enzymes (2, 3), antibodies (4), biologics (5, 6), and peptides (7), these macromolecules navigate an immense range of cellular functions, offering a high degree of specificity and efficacy that traditional small molecule drugs might lack. Capitalizing on our deepening understanding of protein structure and function, protein engineering enables the creation and optimization of therapeutics that can outperform natural molecules in stability, affinity, selectivity, or function. Moreover, it promises the future of precision medicine, with tailored therapies that are designed to align with individual genetic blueprints (8).

In the era of artificial intelligence (AI), the concept of *de novo* design has expanded to include structure-independent methods for rapid sequence generation. Although still in their infancy compared to traditional approaches, these methods hold the potential to revolutionize the field of protein design. The rise of natural language processing (NLP) has significantly advanced protein sequence analysis. Transformer models, which employ self-attention to capture pairwise interactions across the entire input sequence, have surpassed RNNs and LSTMs with their wider reference window. These transformer-based approaches have proven their mettle in machine translation, text summarization, and question answering, as exemplified by popular networks like BERT (9) and GPT (10).

Moreover, transformer-based models pretrained on protein sequences (11–13) have shown competitiveness with other methods by incorporating protein-focused pretraining tasks. This transfer learning approach, often combined with multitask learning, enhances performance and boosts downstream applications for robust protein representation learning. ProSE (14), for instance, applied protein language models to predict structural similarity, secondary structure, residue-residue contacts, and transmembrane regions with promising results. Similarly, ProtGPT2 (15), a variant of GPT-2 (16) trained on a large dataset of protein sequences and associated functional properties, was fine-tuned on specific protein families to generate novel sequences with improved accuracy and novelty compared to existing methods.

The field of protein design has also explored controllable generation, using explicit or implicit control signals to guide generation and meet specific conditions. CTRL (17) is a successful example of this approach in language generation, where a protein language model is conditioned on text properties such as domain and style. In a similar vein, ProGen (18) trains a conditioned pLM on amino acid sequences using conditioning tags representing protein properties. The model, trained on a large dataset of wild-type proteins, generated approximately 1M sequences with 72% of the tested being expressed similarly to natural enzymes. These artificial enzymes exhibited enhanced resilience against small mutations compared to natural proteins.

The advancements in protein language modeling have been nothing short of remarkable, achieving performance levels and a breadth of downstream applications that were unimaginable just a few years ago. However, this rapid progress has been accompanied by a significant rise in computational demands. This emerging pattern, inspired by the so-called scaling laws born from advancements in NLP (19), could overpass the intricate differences distinguishing natural language from the language of proteins.

This quest for larger models has catalyzed a dramatic escalation in the size of pLMs, from a modest one million parameters to an astounding several billion. ESM-1 (20) was a pioneer in this regard, scaling up to 43 million parameters as early as 2019. This was followed by ProtGPT2, RITA (21) and ProGen2 (22) which pushed the envelope further to 15 billion parameters. One of the most recent examples of this trend is xTrimoPGLM (23), which has scaled up to a staggering 100 billion parameters.

However, the assumption that larger models invariably lead to better performance neglects several critical aspects, including the computational burden and specificity for the given task to solve (24). While model size is undoubtedly a significant factor in achieving the objectives of protein language modeling, the escalating computational demands and task-specific optimization often overlooked in the race for scalability, risk elevating the barriers to research innovation, potentially sidelining new perspectives and discoveries. Similarly, the assumption that larger datasets inevitably yield more powerful insights is increasingly being challenged, underscoring the fact that meticulously curated, smaller datasets can offer competitive, if not superior, results (25). Moving forward with protein language modeling, it becomes imperative to take these factors into account, striving for a measured and sustainable approach to model development and application. Such a comprehensive perspective will be critical for appropriately utilizing the full potential of protein language models while maintaining efficiency and accessibility.

In this study, we delve into an efficient approach in protein language modeling by significantly downsizing the UniRef50 dataset (26) and fitting it to a smaller transformer model. This exploration is driven by the hypothesis that smaller, more efficient models can still yield valuable insights, potentially making protein engineering research more accessible and cost-effective. We also investigate the potential of incorporating conditional tags into our model, an approach that could offer more nuanced control over the model’s outputs. We tested our approach on the malate dehydrogenase (MDH) family of enzymes, a widely used group of proteins that has become the standard test in protein generation.

We validate our approach through a combination of bioinformatics analyses and molecular dynamics simulations, ensuring that our generated sequences are not only theoretically sound but also likely to fold correctly and remain stable in real-world conditions, seeing that a vast majority of our generated sequences demonstrate proper folding and stability, underscoring the effectiveness of our approach. Remarkably, despite the reduced size of our model and dataset, we achieve comparable performance in sequence generation to that reported by bigger models. This work, therefore, presents a promising direction for future research in protein engineering, emphasizing efficiency and accessibility without compromising performance.

## Results

### Pretraining on an optimal amount of evolutionary data

Our primary objective was to establish a benchmark for *de novo* protein sequence generation using a small-scale protein language model (SS-pLM). The SS-pLM was pretrained on a curated subset of the Uniref50 dataset, consisting of 1.7 million sequences. This subset offered comprehensive coverage of the enzymatic sequence space and an optimal amount of evolutionary data, enabling the model to learn generic and informative representations encoding the semantic and syntactic information of proteins (27). Further details regarding the dataset acquisition are provided in the methods section.

The evaluation of our model pretraining involved assessing traditional metrics in natural language processing tasks, such as perplexity and accuracy, as detailed in the methods section. Perplexity, measuring the language model’s sequence prediction capability, showed lower values denoting superior performance. Despite the reduced dataset, our SS-pLM achieved perplexities comparable to those of high-quality language models. Perplexity comparisons with uniform and empirical baselines, where amino acids are sampled uniformly and based on observed frequencies, respectively, showcased significant improvements [Table 1]. In terms of accuracy, the model demonstrated a 20% hard accuracy and up to 28% soft accuracy in the corrupted token prediction task, considering substitution frequencies from the BLOSUM62 (28) substitution matrix. This suggests that most incorrect predictions correspond to evolutionary plausible mutations, implying that the model emulates the natural mutagenesis process when proposing new sequences.

**Table 1.** Perplexity (PPL) and accuracy comparisons of the fine-tuned SS-pLM with uniform and empirical baselines. Perplexity measures the predictive performance of the model, with lower perplexity indicating higher quality language models. The hard accuracy evaluates individual amino acid errors, while the soft accuracy, incorporating BLOSUM62 substitution matrix, penalizes predictions based on substitution frequency, considering natural protein substitutions less harshly.

| Model | PPL | Hard acc. | Soft acc. |
| --- | --- | --- | --- |
| Uniform baseline | 12.25 | 4% | 11% |
| Empirical baseline | 11.03 | 6% | 14% |
| SS-pLM | 6.83 | 20% | 28% |

Given the influence of biochemical properties on the interchangeability of amino acids within protein contexts (29), the model is expected to capture these patterns and build a representation space that reflects biochemical knowledge. To further assess the model’s encoding of physicochemical properties, we employed *t*-SNE to project the weight matrices of the initial and final embedding layers into 2D. The projections [Figure 1a] demonstrate that the input embedding space lacks a discernible distribution, while the output embedding space [Figure 1b]exhibits distinct clusters representing hydrophobic and polar residues, aromatic amino acids, and organization by molecular weight and charge. This indicates that the model successfully captures biochemical knowledge and patterns in the amino acid language.

**Fig. 1.**
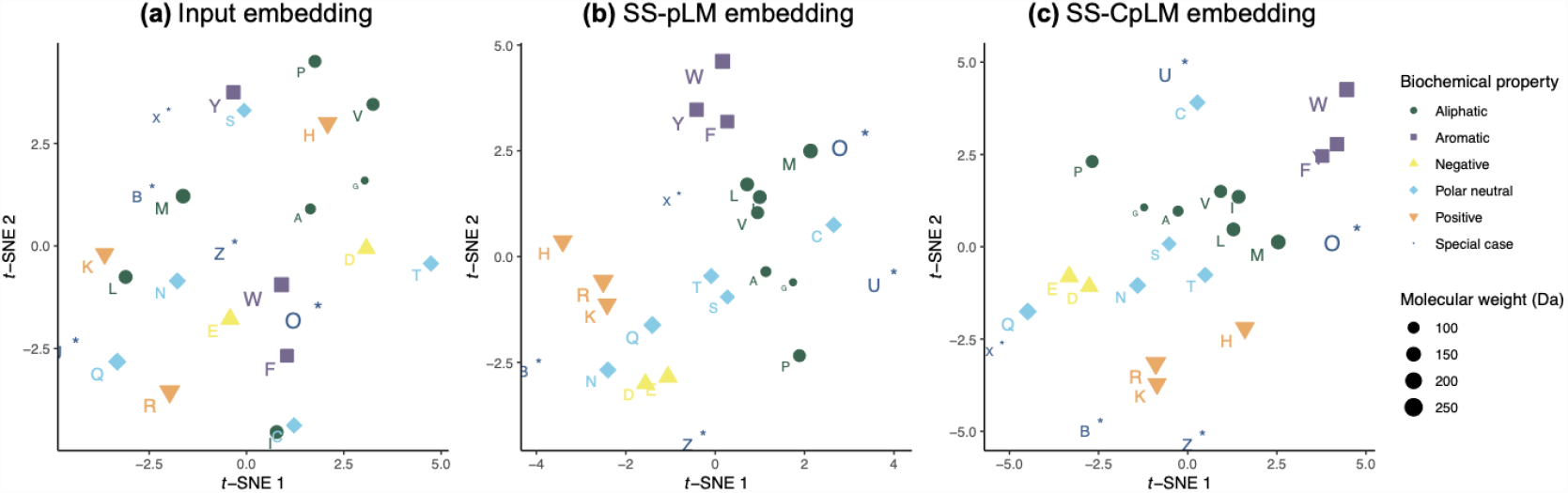
Visualization of amino acid physicochemical properties in pretrained model embeddings using *t*-SNE. The figure depicts the *t*-SNE projection of weight matrices from the final embedding layers of the two pretrained foundation models compared to a random initial embedding layer.

### Incorporation of biological constraints

We wanted to assess whether the incorporation of conditioning tags could yield further improvements in the model’s performance. Following the same architecture and training approach as before, we introduced conditioning tags at the beginning of sequences to represent the first three EC numbers, signifying the enzymatic reactions of each protein in the dataset. Through this approach, we developed a conditional smallscale protein language model (SS-CpLM) that successfully captured the biochemical properties of amino acids in its embeddings [Figure 1c]. Using the same rationale, we evaluated the conditioned foundation model’s ability to capture the distinct groups of enzymatic reactions denoted by conditioning tags and create a corresponding representation space reflecting this chemical knowledge. However, upon analysis, the learned embedding space displayed no discernible distribution [Supplementary Figure 1]. Moreover, we identified a bias towards oxidoreductases in the source dataset, which potentially contributed to the model’s limitations in effectively learning the desired associations in this context.

### Structural and biochemical properties of artificial sequences

To assess protein generation capability within a specific family, we conducted fine-tuning on both SS-pLM and SS-CpLM models using a dataset specific to the MDH family of enzymes encompassing 17,000 bacterial sequences. Subsequently, artificial sequence datasets were generated for each model, and bioinformatic analyses were carried out to validate structural and biochemical properties of the sequences.

At the sequence level, we compared the amino acid content and physicochemical properties of the fine-tuned SS-pLM generated set with natural MDH sequences [Figure 2]. Our analysis involved a comprehensive set of amino acid groups, including Hydrophobic, Hydrophilic, Aromatic, Small, Positive, Negative, Aliphatic, Hydroxyl/Sulfur, Polar Uncharged, and Charged. Statistical tests [Supplementary Table 1] revealed that the model’s fine-tuning successfully aligned the amino acid proportions with those seen in natural proteins. This was remarkable, given the inherent variability in amino acid composition across different protein families. The model maintained a high degree of conservation for most of the amino acid groups, suggesting that it has acquired a sophisticated understanding of biochemical principles. However, three groups - Negative, Aliphatic, and Charged - showed significant differences (p < 0.05) between the model’s generated sequences and natural proteins. This indicates that while the model is generally able to introduce synonym mutations that are biochemically plausible, there are certain physicochemical properties that it does not perfectly match with natural sequences. Despite these differences, the findings provide strong evidence of the model’s capacity to acquire biochemical knowledge and recognize underlying patterns during pretraining and fine-tuning.

**Fig. 2.**
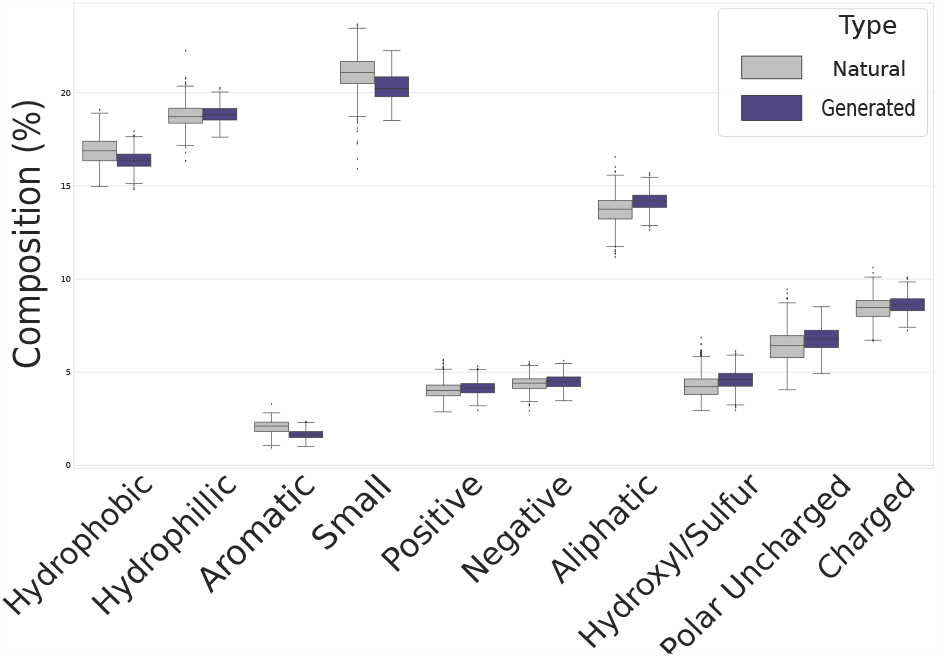
Distribution of amino acid groups of natural and generated sequences. Amino acid percentages for *de novo* and natural sequences were compared by group. Through statistical analysis we concluded that most of the groups were did not have any significant difference.

We then employed multiple sequence alignments to gauge the novelty and resemblance of the fine-tuned SS-pLM generated set of proteins to their natural counterparts. The artificial sequences exhibited an average percent identity of 29% with natural sequences of the enzyme family, spanning up to 89%, and 45% within the generated set. In contrast, random amino acid sequences displayed a mere 9% identity with natural sequences. A higher percent identity signifies closer relatedness between sequences. These compelling findings underscore the model’s capacity to generate a diverse array of sequences that are distantly related to natural proteins, ruling out the possibility of mere random sampling. For further analyses, we honed in on the generated sequences showing percent identities ranging from 60% to 85%, enabling us to scrutinize the model’s ability to produce *de novo* proteins that still retain similarities to those naturally found in biological systems.

In order to simplify the computation of identities and similarities both metrics were calculated using the MDH *E*.*coli* reference sequence (Uniprot ID: P61889) as a benchmark. The distribution pattern unveils that the generated sequences stretch across a diverse range of identities, hitting a minimum of 20% relative to the reference sequence. This variation underscores the model’s adeptness at weaving a substantial number of mutations into the generative process. Yet, the similarities never fell below 80%, implying the model’s discernment in the incorporation of amino acid substitutions [Figure 3]. It adheres to a conservative approach, ensuring that despite the mutations, there is a preserved core resemblance to the reference sequence.

**Fig. 3.**
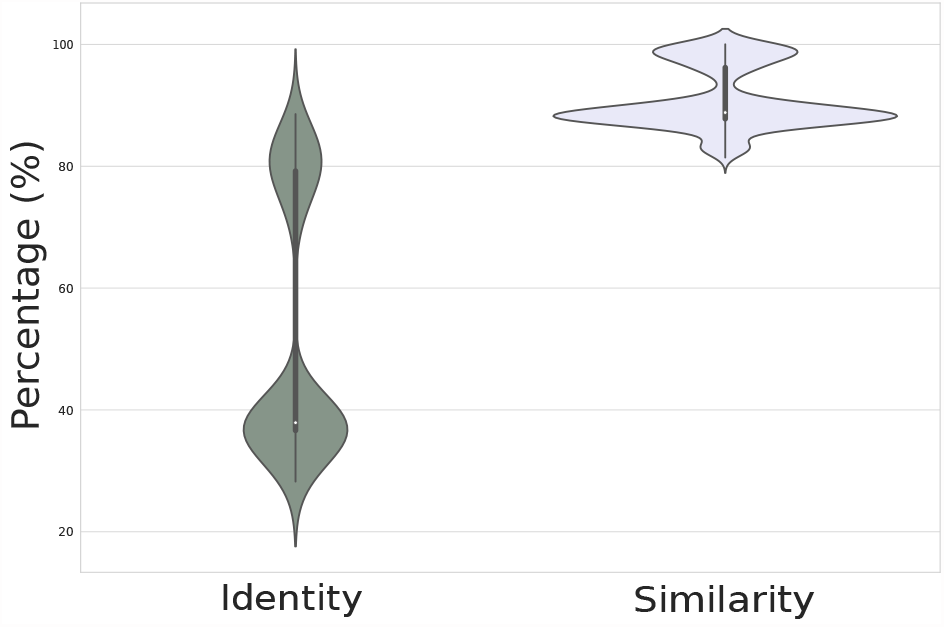
Identity and similarity of generated sequence to reference (*E*.*coli*) The *E*.*coli* MDH sequence was selected to check similarity and identity of generated sequences. Sequences showed a wide range of identities while maintaining a high percentage of similarity.

In our subsequent analysis, we conducted an investigation to ascertain whether the artificial proteins exhibit a degree of similarity in terms of secondary structure element content when compared to natural proteins. To achieve this, we employed PSIPRED predictions to compute the secondary structure content for sequences generated by both the foundation models and the fine-tuned models. Additionally, we included datasets comprising natural and random sequences in this comprehensive examination. Upon inspection of the proportions of secondary structure content [Table 2], we made a notable observation. Specifically, the fine-tuned models displayed an ability to generate sequences with secondary structure contents that bear a strong likeness to those commonly observed in the natural protein space. This finding suggests that pretraining alone may not suffice to generate proteins specific to a particular protein family. The superior performance of the fine-tuned models over the foundation models further supports this point, with the foundation models surprisingly demonstrating even poorer results than random sampling in this context.

**Table 2.** Comparison of secondary structure content between natural, generated and random sequences. The table presents the proportions of secondary structure content computed using PSIPRED predictions (n = 10,000 sequences/dataset).

| Content | Natural | Fine-tuned SS-pLM | Fine-tuned SS-CpLM | SS-pLM | SS-CpLM | Random |
| --- | --- | --- | --- | --- | --- | --- |
| Alpha-helical | 41.55% | 40.39% | 39.74% | 53.19% | 55.37% | 34.75% |
| Beta-sheet | 20.56% | 20.49% | 20.00% | 16.44% | 15.00% | 21.81% |
| Coil | 37.50% | 39.12% | 40.26% | 30.37% | 29.63% | 43.43% |

We next examined the fitness of generated sequences in terms of their ability to fold into stable ordered structures. We evaluated the sequences using ESMFold predictions, considering two key metrics: predicted local Distance Difference Test (plDDT) scores and Predicted Aligned Error (PAE) values. The plDDT scores, ranging from 0 to 100, serve as a measure of confidence and structural order, with higher scores indicating higher confidence. Meanwhile, the PAE graph represents the expected positional error between predicted and actual protein structures, with lower PAE values indicating reliable alignments. Our analysis, [Supplementary Figure 3] revealed that the natural dataset exhibited a significant average plDDT value of 93%, underscoring the high degree of structural coherence observed in naturally occurring proteins. Remarkably, among the artificial sequences [Figure 4], the ones generated using the fine-tuned SS-pLM presented the most elevated plDDT scores and the most diminished PAE values. With an average plDDT score of 87.9%, these sequences showcased the generation of reliable and well-ordered structures. Conversely, the fine-tuned SS-CpLM exhibited less structured and more diverse outcomes, characterized by lower confidence levels and an average plDDT of 70.3%. In contrast, the foundation models themselves produced completely unreliable structures, displaying lower plDDT scores and higher PAE values, with an average plDDT score of 46%. Ultimately, the predictions from the random dataset revealed a markedly lower mean plDDT value of 26%, indicative of a considerable lack of structural order in randomly generated sequences. This outcome reinforces and strengthens the aforementioned results, emphasizing the similarity in structural integrity exhibited by the natural and artificial sequences, in sharp contrast to their random counterparts.

### Assessing the *in silico* viability of generated enzymes

We performed molecular dynamics (MD) simulations of the generated sequences by the SS-pLM to further validate their overall quality and structural integrity. First, we clusterized a sample of 1000 generated MDH sequences at 65% minimal sequence identity using MMseqs2 (30), and then, we took the centroid sequence of each cluster, being a total of 40 sequences. 8 of those 40 sequences did not have the archetypical catalytic histidine residue of MDHs and 2 did not have one of the stabilizing Arg residues in the active site (these last two were still selected for MD validation). The most populated cluster of these sequences was 17, while the other clusters contained less than 10 sequences. Thus, almost all sequences have the critical key residues of the active site. We predicted an accurate structure from the 32 remaining sequences using AlphaFold. In agreement with the results obtained with ESMFold predictions, the AlphaFold models were confident with an average plDDT score of 91.97 % [Supplementary Figure 4]. Only 2 generated enzymes (SG-289 and SG-342) did not have an AlphaFold model with an average plDDT above 90 %.

**Fig. 4.**
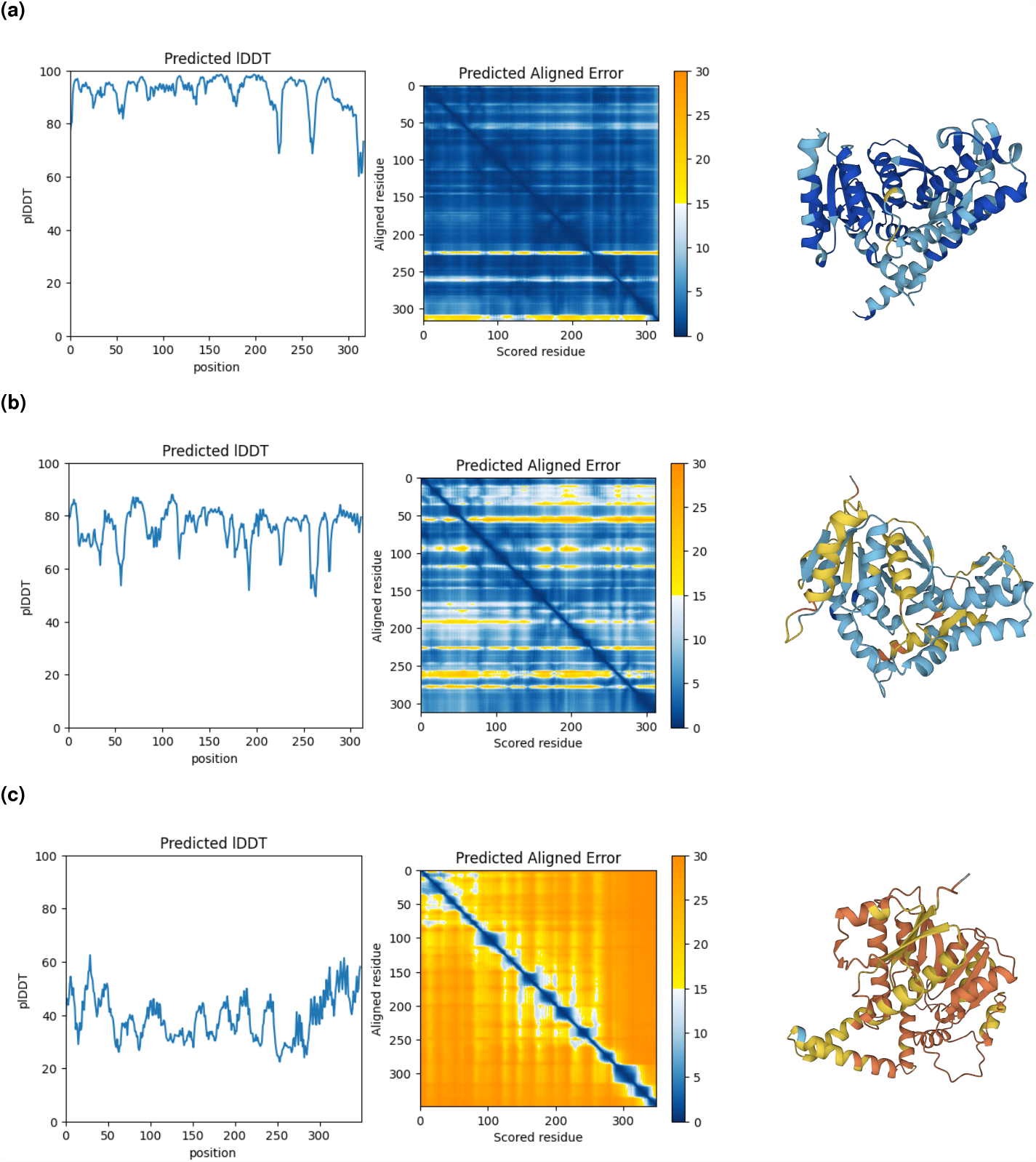
Protein folding analysis of generated proteins using ESMFold predictions. Comparison of pLDDT scores and PAE values for a randomly selected protein sequence generated by different models: (a) fine-tuned SS-pLM, (b) fine-tuned SS-CpLM, and (c) SS-pLM. The predicted structures are shown on the right and are color-coded based on pLDDT values per amino acid location. Blue indicates confident predictions (pLDDT > 0.9), while orange indicates low confidence (pLDDT < 0.5).

The holo MD simulations of the generated MDHs were performed with the best model from the AlphaFold calculation. Likewise, we simulated a generated sequence with a lower mean plDDT score (72.9 %) and two natural enzymes as controls (UniProt ID: [P9WK13, G4T3S9], PDB ID: 5KVV) from the BRENDA database. The analysis of the MD simulations consisted in checking the structural and functional integrity of the generated sequences by examining the distribution of the global root-mean-square deviation (RMSD) of the protein backbone [Supplementary Figure 6], the local RMSD of the active site (catalytic residues and cofactor) [Supplementary Figure 7], and the catalytic distances [Supplementary Figures 5-12]. The overall structure of all the generated sequences seemed to be stable as the natural enzymes except for 7 (SG-249, SG-256, SG-274, SG-733, SG-92, SG-975, and SG-985), while the local geometry of the active site showed a similar behavior to the natural MDHs. All the distances between the substrate and either one catalytic residue of the protein or the cofactor showed very high values because malate exits the active site after ∼ 100ns of simulation. In fact, we measured the fraction of near-attack conformations (NAC) and the vast majority of generated MDHs had some catalytic poses during the simulation [Supplementary Table 2]. Surprisingly, one of the natural enzymes did not show NACs in any of the MD replicas. The main interaction between the catalytic residues of the active site was also preserved in the generated sequences with median values between 2.5 and 7.5 Å. To fully confirm that the generated enzymes preserved the functional fold, we measured the distance between the O atoms connected to one of the P atoms and the H of an amino backbone involved in stabilizing the *NAD*^+^ interaction [Supplementary Figure 13]. All seemed to remain with the cofactor bound to the active site in all MD replicas except for 4 (SG-108, SG-256, SG-975, and SG-985) of which 3 also showed bad global RMSD values, indicating a possible unfolding of the structure. The centroid sequences of the most populated clusters of the sample of 1000 generated sequences (SG-282 cluster with 386 and SG-708 cluster with 352) showed good values in all the measured metrics, meaning that the majority of generated sequences are realistic MDHs.

### Quality of generated sequences under low data regimes

The resilience of the SS-pLM under low data regimes was further examined by reducing the size of the fine-tuning dataset to 10,000, 5,000, and 1,000 sequences from the MDH family. Our goal was to investigate the influence of the training dataset size on the quality and diversity of the sequences generated by the model.

As expected, we observed a decline in the quality of generated sequences with the reduction of the training dataset size. When we compared the generated sequences to their natural counterparts, we found that the sequences from the 10,000, 5,000, and 1,000 fine-tuned models progressively deviated more from the natural MDH sequences [Supplementary Figure 2]. This deviation was characterized by a lower overall percentage of identity and lower similarity, indicating that the model was more likely to introduce non-similar substitutions, which could potentially impact the folding or the catalytic machinery of the enzyme.

Alongside this decline in sequence similarity, we also observed a decrease in the diversity of the generated sequences as the dataset size was reduced. This was evidenced by a higher prevalence of identical or highly similar sequences within the generated datasets, particularly for the model finetuned on 1,000 sequences. This trend suggests that the model, when trained on smaller datasets, tends to generate a narrower range of sequences, potentially limiting its utility in applications requiring a broad diversity of novel sequences.

Interestingly, despite the observed decrease in sequence similarity and diversity, the folding potential of the generated sequences remained relatively stable across all dataset sizes. We used ESMFold to fold the generated sequences and found that the sequences from the 10,000 and 5,000 fine-tuned models demonstrated good folding potentials, with mean plDDT scores of 87.81 and 85.18 [Supplementary Figure 3], respectively, indicating that viable sequence generation is possible even within low data regimes. The 1,000 fine-tuned model presented a more significant shift towards dubious folding with a mean plDDT score of 71.62, but still yielded a percentage of folded sequences within the range of usable enzymes. This result suggests that the model retained its ability to generate structurally viable sequences even under significant data limitations.

## Discussion

Our study has presented a compelling exploration into the capabilities of SS-pLMs in *de novo* protein sequence generation. By focusing on efficiency and accessibility, we have provided a foundation for researchers operating under data and computational constraints, demonstrating that significant progress can be achieved without enormous datasets or high-scale computational resources.

We have elucidated that pretraining SS-pLMs on large datasets of natural proteins allows these compact models to comprehend complex biochemical knowledge and discern intricate patterns within the language of amino acids. This learning, in turn, empowers the models to generate artificial sequences that mirror natural proteins in terms of their structural and biochemical properties. This finding adds credence to the idea that proficiency in protein generation isn’t solely tied to the scale of the model but can be achieved through well-designed learning paradigms on adequately large and representative datasets.

Our study highlights the added benefits of fine-tuning the SS-pLMs on datasets specific to particular protein families. This process not only enhances the quality of generated sequences but also diversifies the protein properties that the model can emulate. Therefore, it amplifies the potential of SS-pLMs to generate proteins with desired properties, such as heightened enzymatic activity or increased binding affinity, thereby opening new avenues for targeted protein design.

The experiments conducted in this study showcase the successful generation of diverse and biologically meaningful protein sequences by our pretrained SS-pLM. The generated samples exhibit confidently predicted structures, as validated by ESMFold predictions. Additionally, these sequences demonstrate significant similarity to natural proteins in terms of both structural and functional characteristics. This remarkable resemblance indicates that the model has adeptly captured essential features of natural proteins during its evolutionary pretraining.

Moreover, we delved into the robustness of SS-pLMs in low data regimes. Our findings suggest that these models maintain the capability to generate high-quality sequences even when the fine-tuning dataset is small. However, we noted a decrease in the diversity of the generated sequences, indicating a trade-off between data size and sequence diversity, a factor that researchers should consider when working under data-constrained environments.

These findings, while promising, are not without limitations. Our exploration focused on a limited number of protein families, raising the question of whether these results would translate across different protein families. Furthermore, we used a limited number of fine-tuning datasets, leaving open the possibility that other datasets might yield differing outcomes.

Despite these limitations, our study underscores the potential of SS-pLMs in making more accessible *de novo* protein sequence generation. By demonstrating that these scaled-down models can yield a diverse array of high-quality protein sequences, we have opened a new alternative in protein sequence generation. Our insights into the factors influencing sequence quality and diversity provide valuable information that can guide future design and refinement of SS-pLMs. With continued exploration and improvement, we believe that this approach can significantly advance protein engineering, facilitating the design of proteins with a wide range of tailored properties.

## Methods

### Data collection

The protein language model underwent initial training on a comprehensive dataset encompassing the entire protein sequence space to capture general features and patterns. We utilized a subset of the UniRef50 dataset (26), retrieved in January 2023, containing approximately 59 million non-redundant sequences grouped into protein clusters sharing at least 50% sequence identity. To reduce the dataset’s size, we retained representative sequences from clusters with more than one member, resulting in a dataset of 22 million sequences. In preparation for future conditional generation objectives, we exclusively retained UniProt entries classified using the Enzyme Commission (EC) system (31), a numerical classification system that categorizes enzymes based on their specific catalytic reactions. This filtering procedure resulted in a dataset of 1.75 million sequences, each associated with a first, second, and third EC number. To promote generalization, we excluded the fourth EC number, which pertains to specific reactions.

Following pretraining, the pretrained model underwent fine-tuning on a specific dataset to optimize performance and tailor it to a specific protein family. Bacterial MDH sequences were retrieved from UniProt (32). Sequences exceeding 512 amino acids were excluded, resulting in a dataset comprising 17,000 sequences. Among these, 300 sequences were randomly chosen for validation, while the remaining 16,700 sequences were used for training.

In both datasets, sequences were arranged into batches and standardized to have the same length in order to streamline the workflow of the implemented models. The sequence length was calculated as the mode of all training sequence lengths plus one standard deviation. Additionally, a dataset of random sequences was created through random sampling from the amino acid alphabet for testing purposes.

### Sequence embeddings and conditioning tags

Before feeding amino acid sequences to a neural network model, they must first be numerically encoded through an encoding scheme that assigns a numerical representation to each amino acid, resulting in amino acid sequences as vector representations or sequence embeddings. To achieve this, protein sequences were encoded using integer tokens, including 28 unique tokens that represent the 20 standard amino acids plus 4 other IUPAC codes such as “B”, “U”, “O”, and “Z”, an unspecified amino acid “X”, a padding token “PAD”, and 2 extra tokens “START” and “END”.

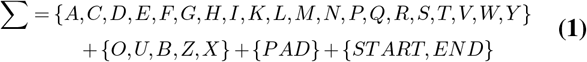

As the implemented architecture requires a fixed sequence length, we randomly selected a subsequence of a protein when encoding sequences longer than the chosen length. To pad shorter sequences, the “PAD” token was added. Additionally, “START” and “END” tokens were added to each sequence, preceding the first amino acid and following the last amino acid, respectively, to assist the language model in processing proteins that exceed the selected sequence length. When exploring conditional generation, we incorporated 340 extra tokens denoting the different combinations of the first, second, and third EC numbers, which were used as conditioning tags. An additional token was introduced to indicate no specific condition, enabling unconditional generation.

### Model architecture

Inspired by the Transformer architecture (33), our basic language model encompasses an encoder and a decoder. The encoder is responsible for transforming input sequences into a continuous representation that retains knowledge and aids the decoder in attending to relevant positions during sequence generation. A crucial component of the Transformer is the self-attention mechanism employed in the multi-headed attention layer. This mechanism computes attention weights for the input and produces an output vector that encodes how each position in the sequence contributes to the representation of other positions. By attending to different parts of the input sequence, the model can capture dependencies and relationships between elements effectively. Another approach used to preserve the order of the input sequence is positional encoding. This positional encoding is applied to the initial embeddings and provides information about the position of each element in the sequence, helping the model distinguish between different positions.

The decoder operates in an autoregressive manner, generating sequences one element at a time using attention information from both the encoder and the previously generated output sequence as inputs. To ensure that the decoder attends only to tokens preceding the current position during sequence generation, a look-ahead mask is applied. This mask prevents the decoder from attending to future tokens, facilitating the modeling of dependencies and ensuring the generation process follows the correct order.

In autoregressive models, the total probability of a sequence *A* = (*a*_1_…*a*_*n*_) is the combination of the individual probabilities for each amino acid given its previous tokens. This distribution can be factorized into a product of conditional probabilities using the chain rule of probability (34), being expressed as:

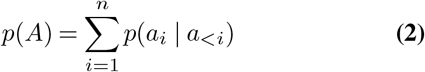

During the training process, the language model with parameters *θ* is trained to generate artificial sequences. This is achieved by minimizing the loss function, specifically the cross-entropy loss, which aims to minimize the negative log-likelihood over the entire dataset *D* = (*A*_1_…*A*_*m*_):

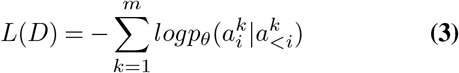

Furthermore, the model is optimized using techniques such as backpropagation and gradient descent to update the model parameters and improve its ability to generate coherent and contextually appropriate sequences.

### Training details

We followed a pretraining and fine-tuning approach [Figure 5] to obtain robust language model’s representations and assess the impact of pretraining on protein generation tasks, including conditional generation. Details of the obtained models, achieved through systematic experiments and performance evaluations under various hyperparameter configurations, are reported [Table 3]. All models utilized AdamW (β1 = 0.9, β2 = 0.999) (35) for optimization, and to monitor their performance, along with training losses, they were evaluated at the end of each epoch.

**Table 3.**
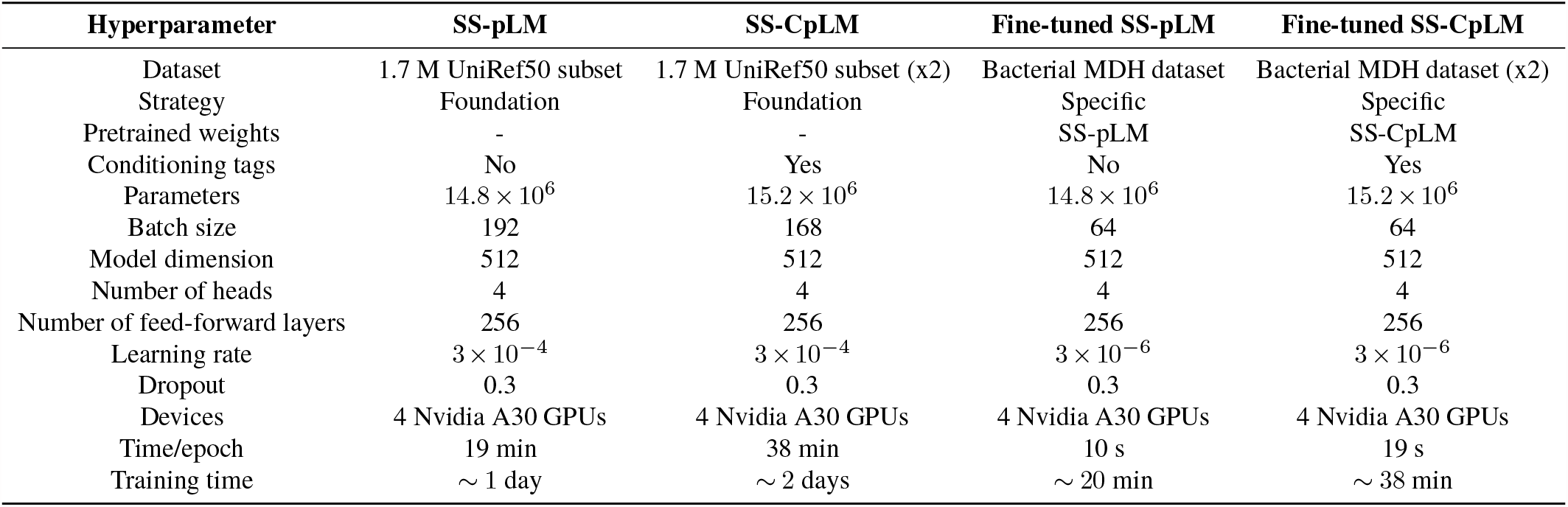
Trained models overview. This table provides detailed information on each model trained throughout the project. It includes whether the model is a foundation model or fine-tuned with pretrained weights, the source dataset used for training, the presence of conditioning tags in the sequences, and the corresponding hyperparameters. The following abbreviations are used: SS-pLM, Small-Scale Protein Language Model; SS-CpLM, Conditional Small-Scale Protein Language Model.

**Fig. 5.**
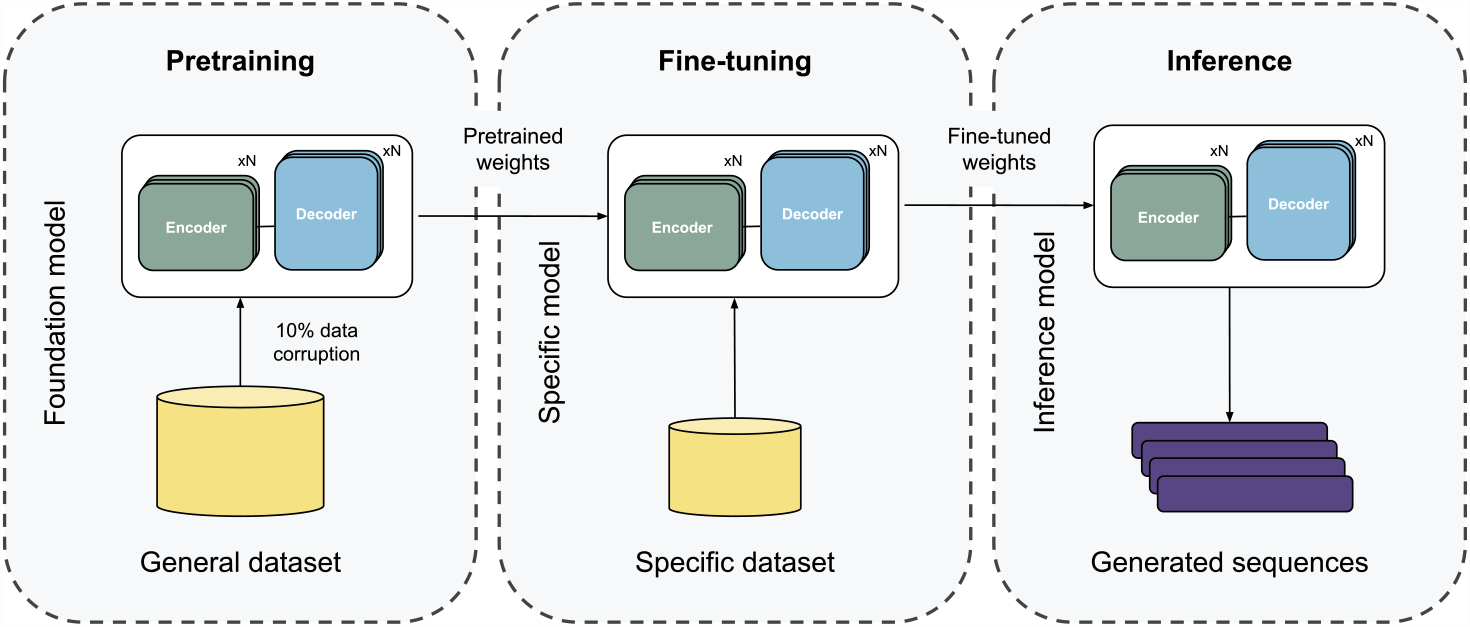
General scheme of the transfer learning-based pretraining and fine-tuning approach. This approach leverages knowledge and representations learned from a large dataset to enhance performance on a specific task with a smaller, specific dataset. Initially, a neural network is trained by randomly initializing the weights and optimizing them to minimize task-related errors. Upon achieving satisfactory training results, the network weights are saved. To train a new network for a different task and dataset, instead of starting from random initialization, the saved weights from the previous network are used as the initial values. In this initialization process, the first network is the pre-trained network, and the second network undergoes fine-tuning.

We pretrained two foundation models with the same architecture using the reduced UniRef50 dataset: a small-scale protein language model (SS-pLM), and a conditional variant of the model with conditioning tags added to sequences, referred to as the conditional small-scale protein language model (SS-CpLM). The causal language model approach with data corruption was employed, where 10% of the amino acids in each sequence were randomly replaced with amino acids drawn uniformly from the real distribution. The models were trained on the next token prediction task, with the loss computed only for the corrupted positions. This objective aimed to impart the models with the intricate language patterns and rules specific to protein sequences corresponding to each combination of first, second, and third EC numbers. To assess the impact of pretraining on protein generation, both SS-pLM and SS-CpLM networks underwent fine-tuning using the bacterial MDH dataset.

For conditioned training, each sequence was included twice: once prepended with the corresponding conditioning tag and once with a tag encompassing all tags. This allowed the model to generate proteins using only sequence data even when no enzymatic function was known.

To examine the impact of reducing the fine-tuning dataset size, we performed fine-tuning on the SS-pLM model using three smaller datasets containing 10,000, 5,000, and 1,000 natural sequences, randomly sampled from the complete bacterial MDH dataset. The training process employed the same architecture, learning rate of 3 × 10^*−*5^, and batch sizes of 64, 32, and 16, respectively.

### Optimization details

We utilized DeepSpeed (36) training strategy and FP16 mixed precision training for parallel computing and accelerated training. Memory usage was reduced by implementing ZeRO stage 2 (37) with optimizer states and gradient offloading. Efficient data processing techniques were applied, optimizing the number of loader worker processes and enabling automatic memory pinning.

### Inference details

To generate coherent sequences with the trained models, we opted to use the beam search algorithm. This algorithm selects the top-k most probable candidates at each time step and discards the rest. The candidate with the highest probability at the end of decoding becomes the output. The beam width, denoted as *k*, is a critical parameter in this process. We experimented with various beam sizes and determined *k* = 3 as optimal. However, sequences generated using beam search often suffer from repetitiveness. To address this, we introduced a repetition penalty to discourage the generation of tokens already present earlier in the sequence.

### Model evaluation

To assess the performance of our pLMs, we employed standard metrics from natural language processing tasks, including perplexity and accuracy. Specifically, we utilized hard and soft accuracy metrics adapted for protein sequence generation tasks, as introduced in ProGen. The hard accuracy assesses each amino acid error strictly, while the soft accuracy incorporates BLOSUM62 (28), a widely used amino acid substitution matrix. The soft accuracy penalizes incorrect predictions based on the substitution frequency in the matrix, giving less penalty for substitutions more likely to occur in natural proteins.

To study the learned properties of amino acids in the protein language, as well as the combinations of EC numbers and their major groups in the conditioning tags, we applied dimensionality reduction using the *t*-SNE algorithm (38). This was performed on the weight matrix of the output embeddings from both foundation models, and one initial embedding used for comparison.

### Sequence evaluation

We created three distinct datasets to assess the quality of generated proteins. First, we obtained a set of 10,000 artificial sequences from each trained model. Second, a natural dataset of 10,000 sequences was randomly sampled from the bacterial MDH dataset. Lastly, we generated a dataset of random sequences with the same length as the previous datasets by random sampling from the amino acid alphabet for testing purposes. Comparative analyses were performed on these datasets using various methods. Artificial sequences were compared to natural ones using BLAST (39). Secondary structure prediction was conducted using PSIPRED (40), considering residues with confidence scores above 0.5 and incorporating PSI-BLAST (39) to enhance prediction accuracy with MSAs from the UniRef50 database. For 3D fold analysis, ESMFold (41) was employed, using the ESM2 protein language model to predict 3D atom coordinates.

### *In silico* protein preparation

The structures of the *de novo* generated sequences were obtained with version 2.3.0 of AlphaFold (42). The *NAD*^+^ cofactor was added from the closest reference found in the PDB hits from AlphaFold. Then, the structures were prepared and protonated at pH 7.0 using Protein Preparation Wizard (43) and PROPKA (44), including fixing side-chains and missing loops using Prime (45), as well as a restrained minimization of the system. The catalytic His residue was constrained to be δ-protonated during the preparation. Before the molecular dynamics (MD) simulations, malate was docked at the active site of the *de novo* generated enzymes using Glide (46). First, the grid of each protein was created with the center being located at the basic N atom of the catalytic His residue in the active site, where the inner box was limited to a cube with an edge of 10 Å. The ligand was sampled as flexible and standard precision was used. 100 poses were extracted and all of them were minimized after the molecular docking with the OPLS2005 force field (47). The best catalytic position for MD simulation was selected based on the post with the best docking score that fulfilled the catalytic distances [Supplementary Figure 5, Supplementary Table 2], representing a near-attack conformation (NAC).

### Molecular dynamics

Four replicas of 500 ns of holo MD simulations were performed with OPENMM (48) to analyze the structural integrity of the active site and the overall structure of the *de novo* generated sequences. A water cubic box (distance of 8 Å between the closest protein atom and the edge of the box) was created around the system using the TIP3P water model, and the system’s charge was stabilized using monovalent ions (*Na*^+^ and *Cl*^*−*^). The protein part of the system was parameterized with the AMBER99SB force field (49). The *NAD*^+^ cofactor parameters were extracted from the AMBER parameter database from the Bryce group (http://amber.manchester.ac.uk/). Malate was parameterized using AmberTools and the GAFF force field (50). The Andersen thermostat and Monte Carlo barostat were applied to the NPT ensemble (constant pressure and temperature of 1 bar and 300 K, respectively). The NVT equilibration lasted 1 ns, and a constraint of 10 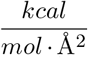 was applied to the system, while the NPT equilibration lasted 2 ns, and a milder constraint of 5 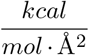 was used. The Verlet integrator, which has a 2 fs time step, was used with constraints between H and heavy atoms. For the non-bonded long-range interactions, a radius of 8 Å was used.

### Future work

As this research is still ongoing, the authors have several significant milestones and implementations yet to be accomplished. One of the key objectives includes a more comprehensive examination of the extent of fine-tuning in terms of sequence space and enzyme quality. This will provide a deeper understanding of the model’s capabilities and its potential applications in protein engineering.

Furthermore, the validation of the methodology is expected to be expanded to encompass other protein families. This extension will not only test the versatility of our approach but also provide a broader basis for its applicability across different protein classes. Crucially, this expanded validation will be accompanied by rigorous experimental verification, ensuring the real-world relevance and effectiveness of our methodology.

In addition to these experimental validations, the authors also explore modifications in the training-generation process. The aim is to leverage the learned protein language to enhance sequence properties deliberately. This could potentially allow for the directed design of proteins with desired characteristics, further extending the utility of our approach.

We also plan to integrate FlashAttention (51), an efficient variant of self-attention, which promises to improve the model’s ability to capture complex relationships within protein sequences. Additionally, we aim to incorporate qLoRA (52), a memory-saving fine-tuning approach, to significantly reduce the computational resources required for model finetuning. This will provide an extra layer of computational efficiency, lowering the entry barrier to our approach.

## ACKNOWLEDGEMENTS

The authors want to thank all Nostrum Biodiscovery team for the helpful everyday discussions and their involvement in supporting in-house research. Special thanks to the IT Department for their quick and invaluable support with the internal cluster.

## Supplementary Material

**Supp. Fig. 1.**
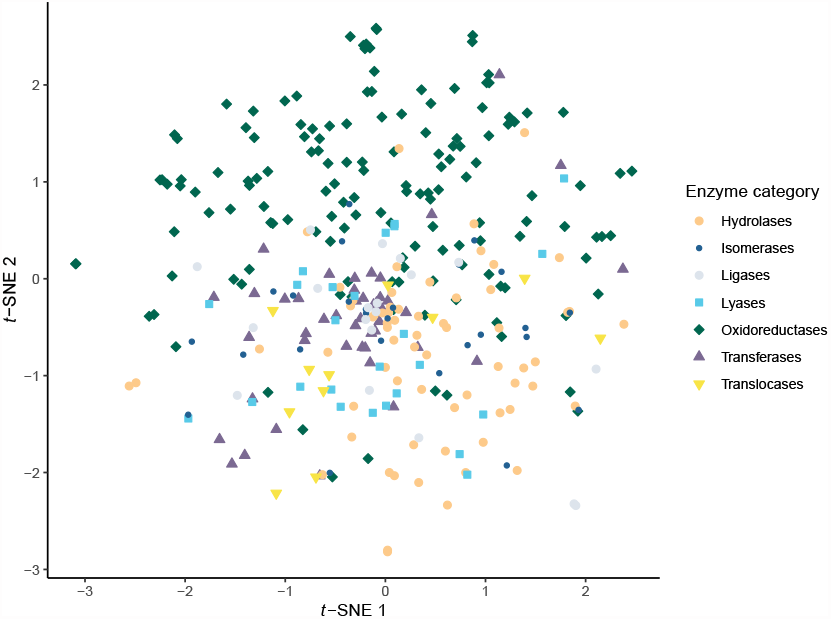
Visualization of amino acid physicochemical properties in pretrained model embeddings using *t*-SNE. The figure depicts the *t*-SNE projection of weight matrices from the final embedding layers of the two pretrained foundation models compared to a random initial embedding layer.

**Supp. Fig. 2.**
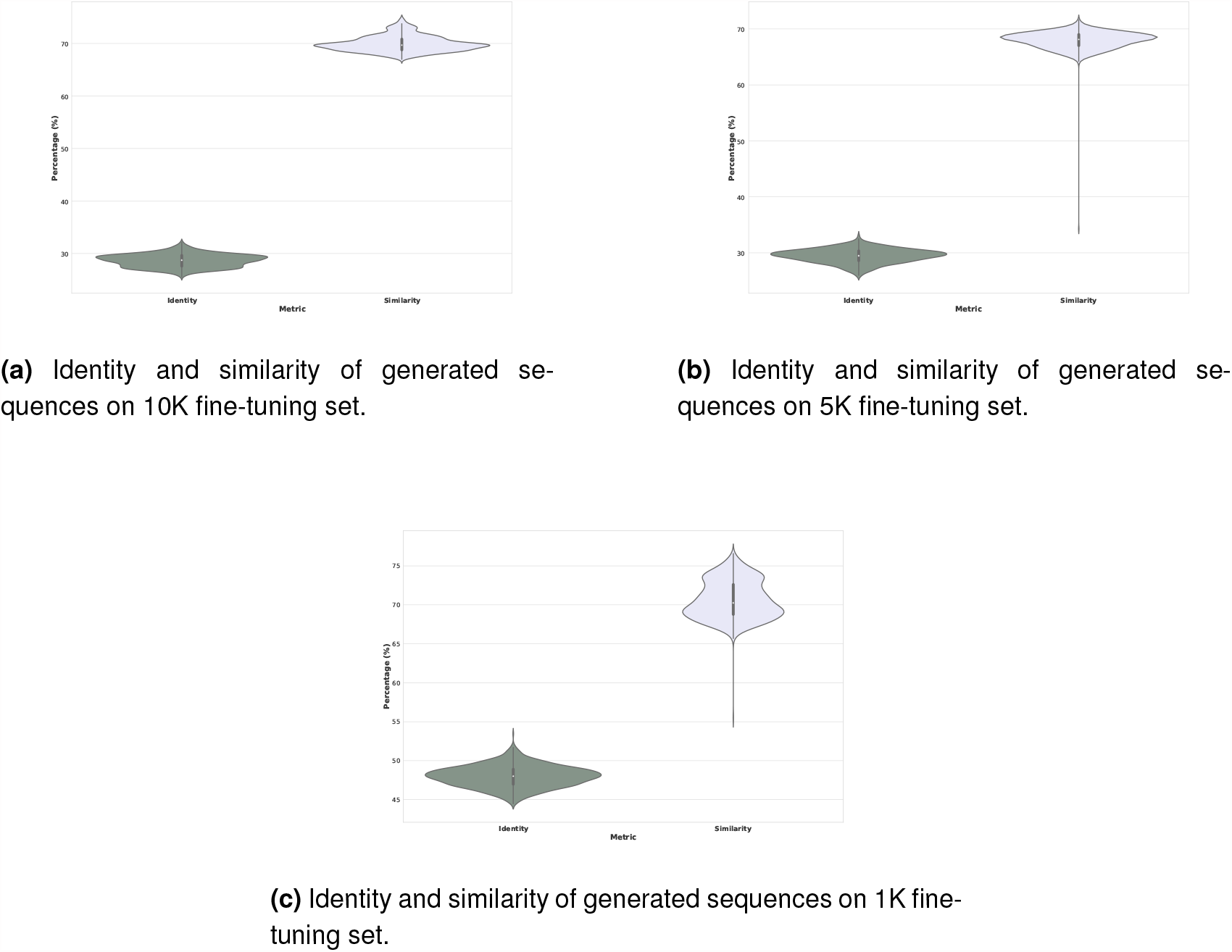
Compilation of identities and similarities of different fine-tuning experiments upon the *E*.*coli* reference sequence. Both metrics were computed for generated sequences out of 10K, 5K, and 1K fine-tunings. Even though sequences still conserve more similarity than identity, we can observe a diminished sequence diversity and an overall lower sequence similarity.

**Supp. Table 1.** Statistical tests amino acid composition. Chi-square statistics and p-values for aminoacid groups. The table presents the chi-square values and the corresponding p-values for various groups including Hydrophobic, Hydrophilic, Aromatic, Small, Positive, Negative, Aliphatic, Hydroxyl/Sulfur, Polar Uncharged, and Charged.

| Group | Chi-square | p-value |
| --- | --- | --- |
| Hydrophobic | 486951.49 | 0.113 |
| Hydrophilic | 466094.17 | 0.865 |
| Aromatic | 275722.52 | 0.080 |
| Small | 545652.78 | 0.214 |
| Positive | 374507.61 | 1.000 |
| Negative | 378590.81 | 0.014 |
| Aliphatic | 500252.49 | 0.006 |
| Hydroxyl/Sulfur | 436724.52 | 0.511 |
| Polar Uncharged | 563271.63 | 1.000 |
| Charged | 469183.73 | 0.002 |

**Supp. Fig. 3.**
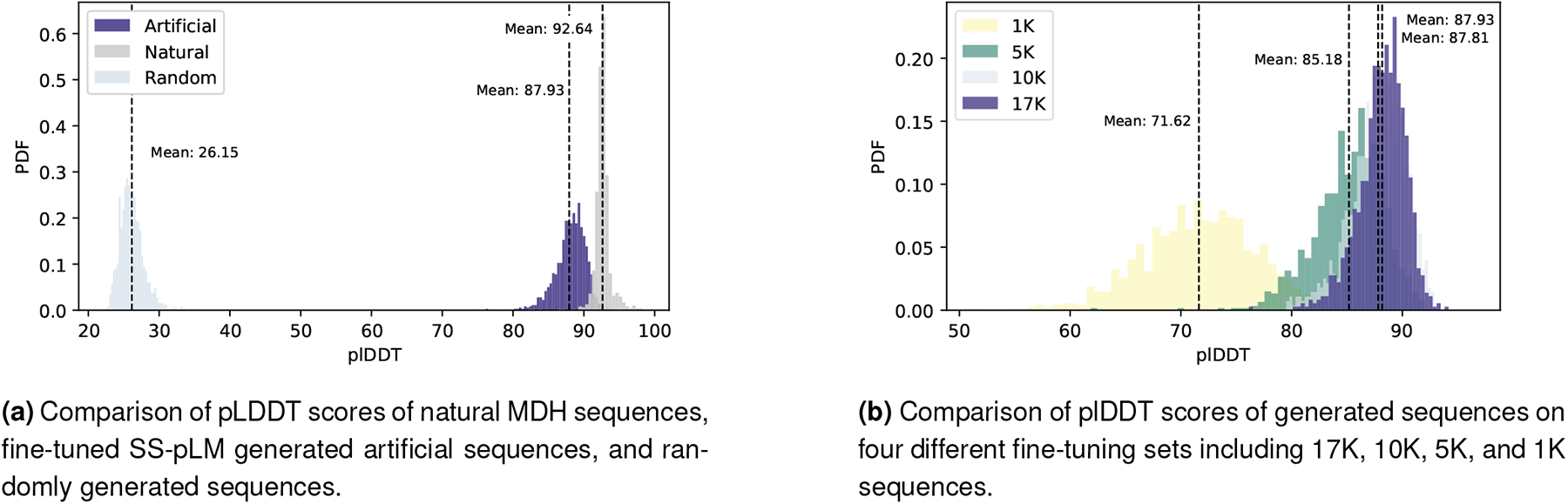
Distribution of pLDDT scores from ESMFold predictions on the distinct protein sequence datasets.

**Supp. Fig. 4.**
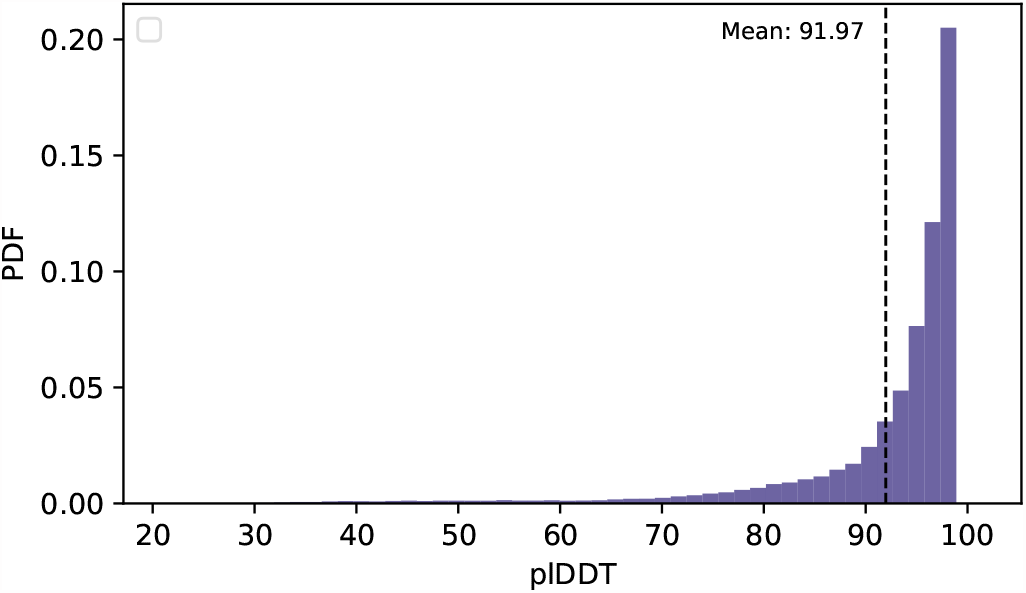
Distribution of plDDT scores of the AlphaFold structures from the sample simulated with MD.

**Supp. Fig. 5.**
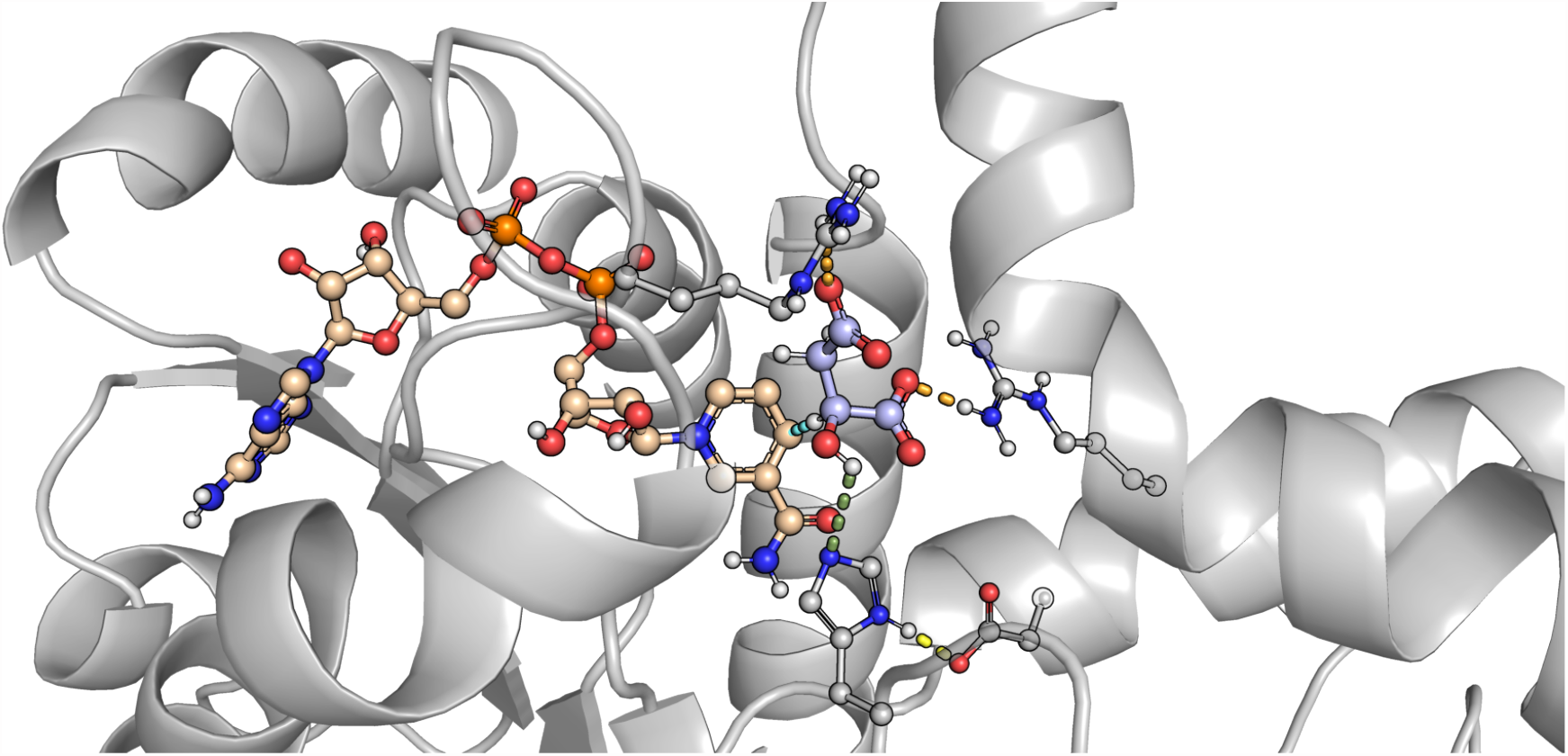
3D representation of a NAC of a simulated MDH with the key distances highlighted. The distance between the *H*^*−*^-accepting C atom of the *NAD*^+^ cofactor and the released H atom of the substrate (named “Cofactor-Substrate distance”) is colored in aquamarine, the one between proton of malate and the basic N atom of the catalytic His residue (named “Histidine-Substrate distance”) in smudge, the distance between the δ proton of the catalytic His residue and one of the carboxylic O atoms of the catalytic Asp residue (named “Aspartate-Histidine distance”) in yellow, and the carboxylic O atoms of the substrate and the H atoms of the basic moiety of the stabilizing Arg residues from the binding site (named “Arginine-Substrate distance”) in orange.

**Supp. Fig. 6.**
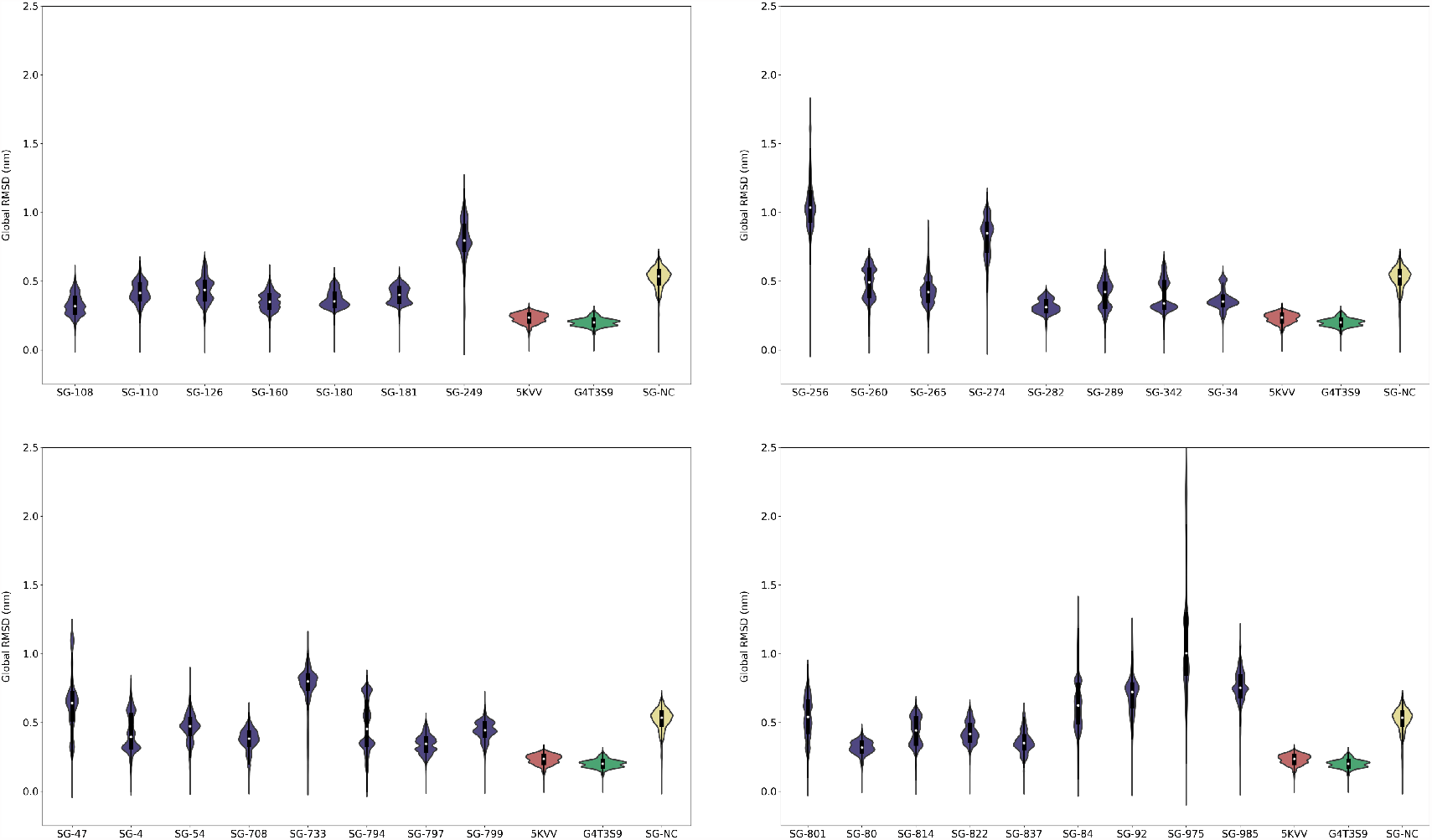
Distribution of the protein’s backbone RMSD of the simulated MDHs. The generated sequences are colored in darkslateblue, while the 5KVV control is in red, the G4T3S9 control is in green, and the generated sequence with bad plDDT in the ESMFold prediction is in khaki.

**Supp. Fig. 7.**
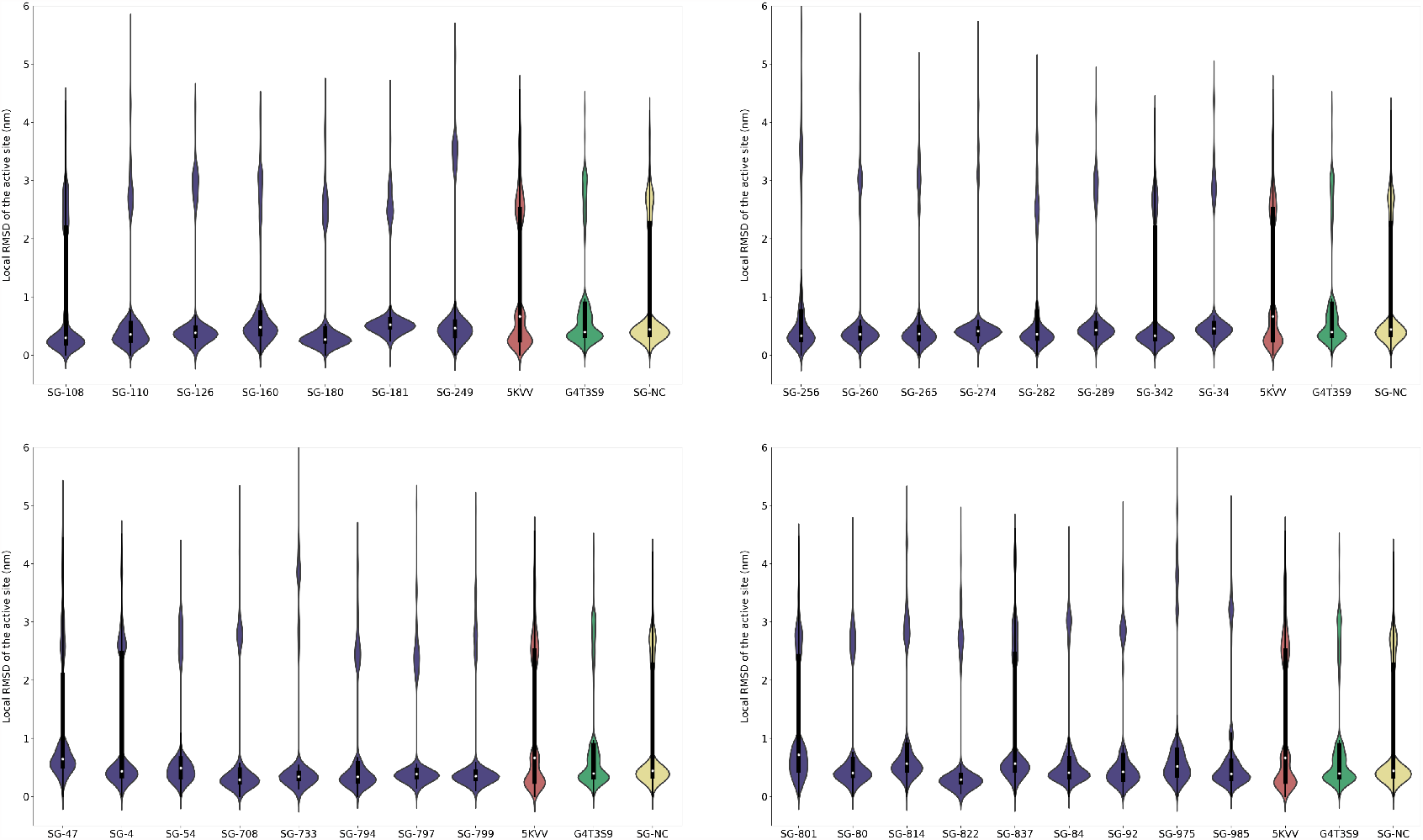
Distribution of the active site’s RMSD of the simulated MDHs. The generated sequences are colored in darkslateblue, while the 5KVV control is in red, the G4T3S9 control is in green, and the generated sequence with bad plDDT in the ESMFold prediction is in khaki.

**Supp. Fig. 8.**
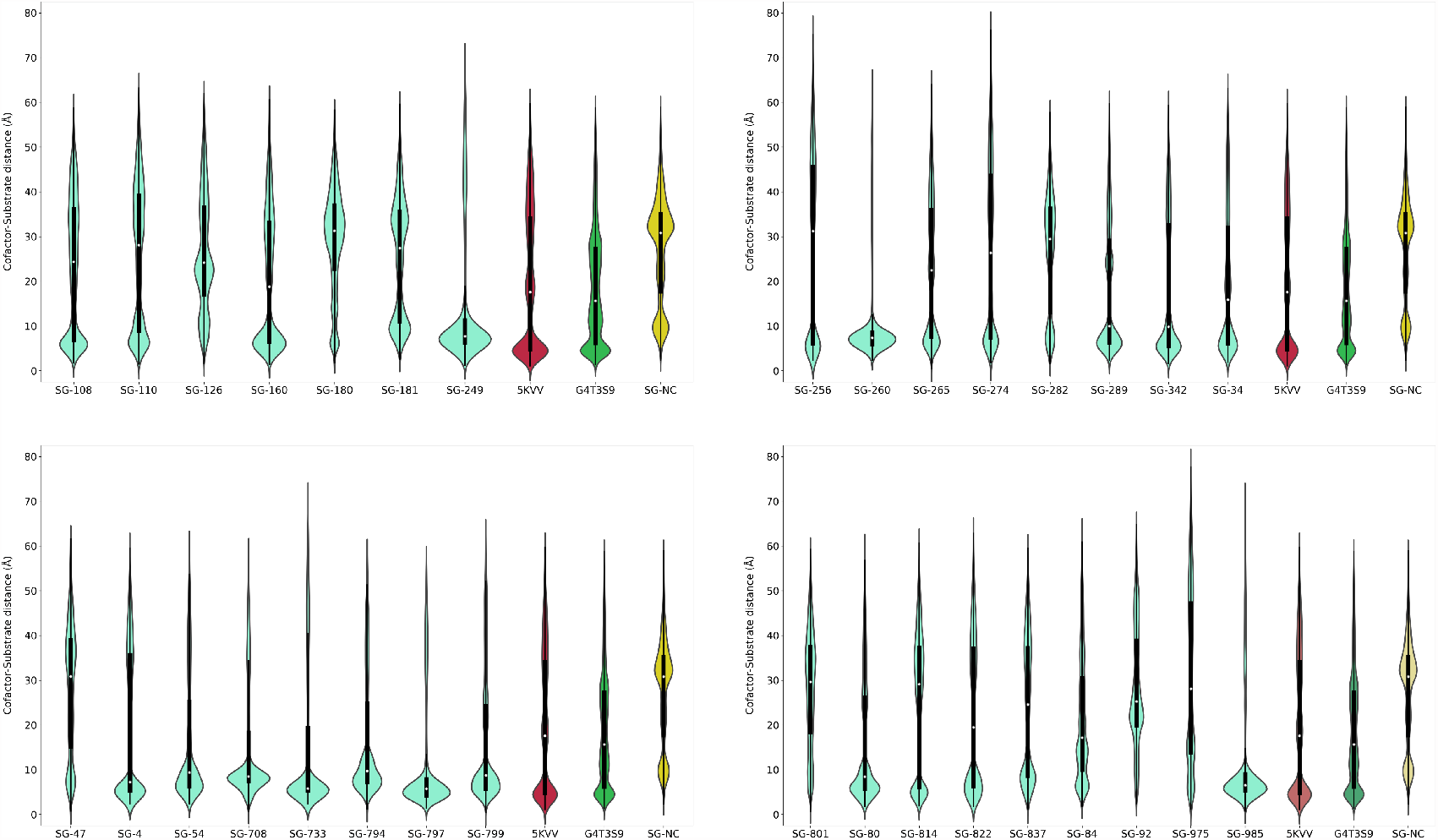
Distribution of the Cofactor-Substrate 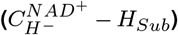 distance of the simulated MDHs. The generated sequences are colored in aquamarine, while the 5KVV control is in red, the G4T3S9 control is in green, and the generated sequence with bad plDDT in the ESMFold prediction is in khaki.

**Supp. Fig. 9.**
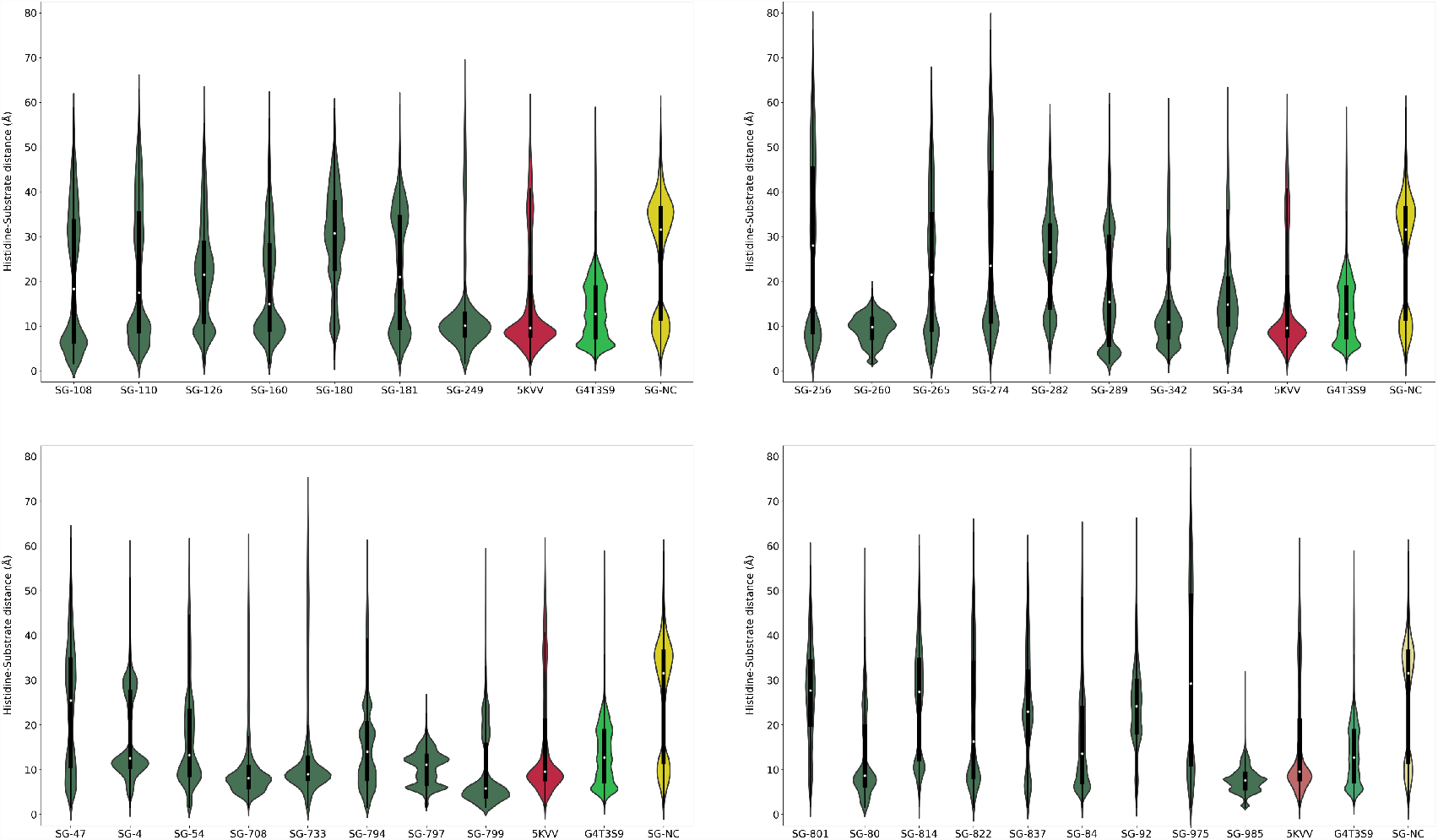
Distribution of the Histidine-Substrate 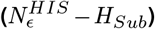 distance of the simulated MDHs. The generated sequences are colored in smudge, while the 5KVV control is in red, the G4T3S9 control is in green, and the generated sequence with bad plDDT in the ESMFold prediction is in khaki.

**Supp. Fig. 10.**
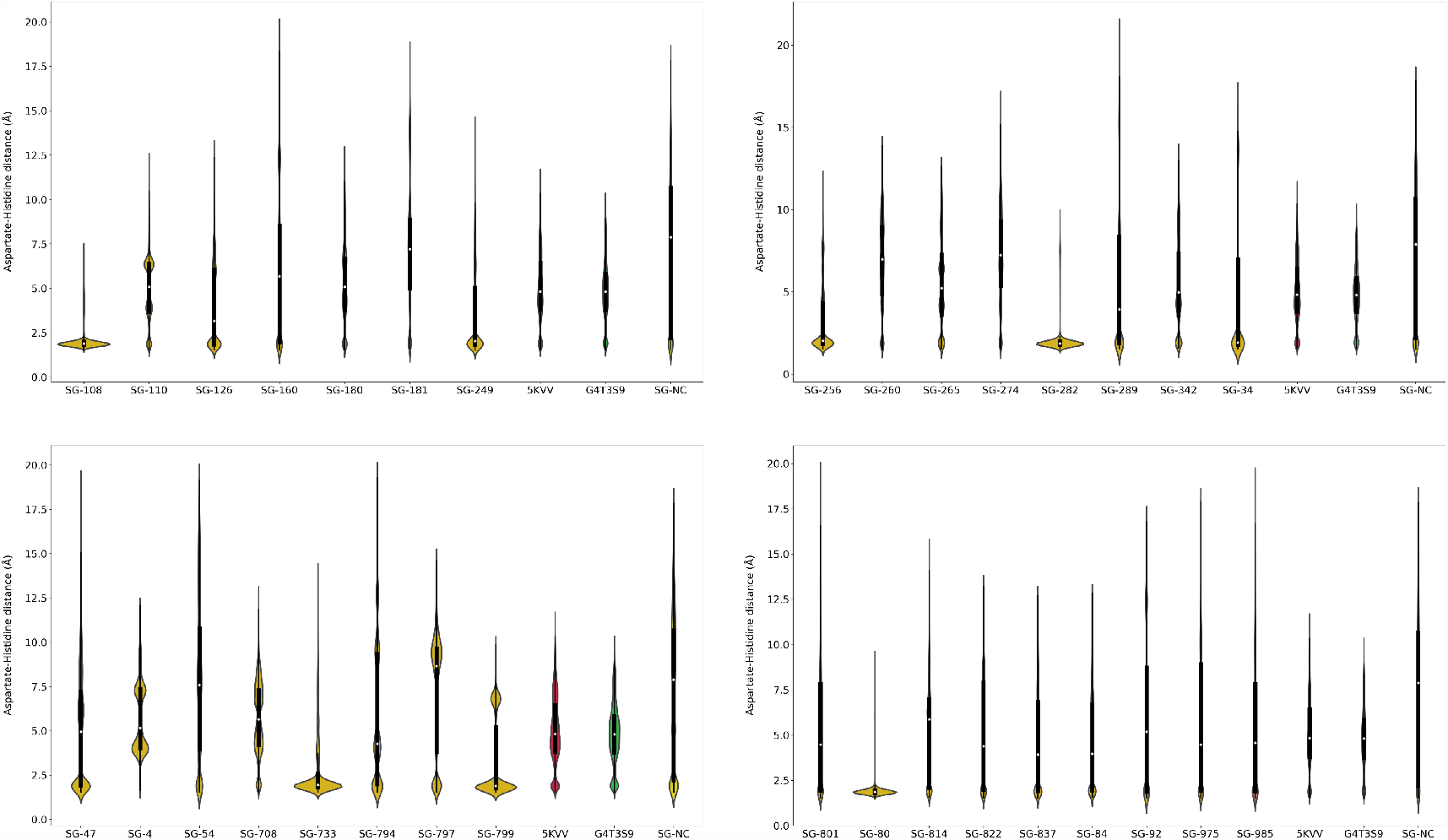
Distribution of the Aspartate-Histidine 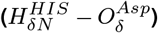 distance of the simulated MDHs. The generated sequences are colored in yellow, while the 5KVV control is in red, the G4T3S9 control is in green, and the generated sequence with bad plDDT in the ESMFold prediction is in khaki.

**Supp. Fig. 11.**
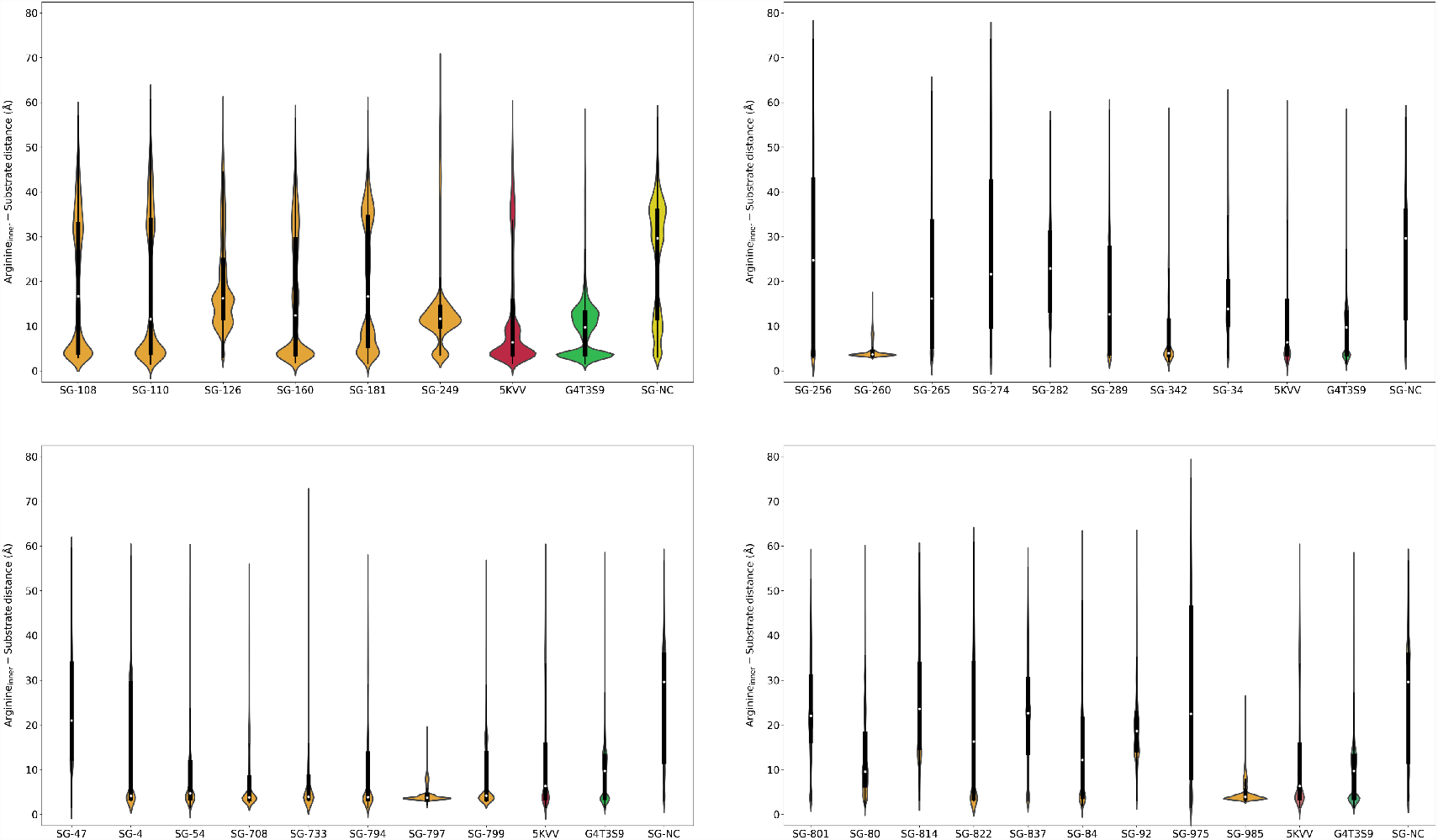
Distribution of the inner Arginine-Substrate 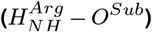 distance of the simulated MDHs. The generated sequences are colored in orange, while the 5KVV control is in red, the G4T3S9 control is in green, and the generated sequence with bad plDDT in the ESMFold prediction is in khaki.

**Supp. Fig. 12.**
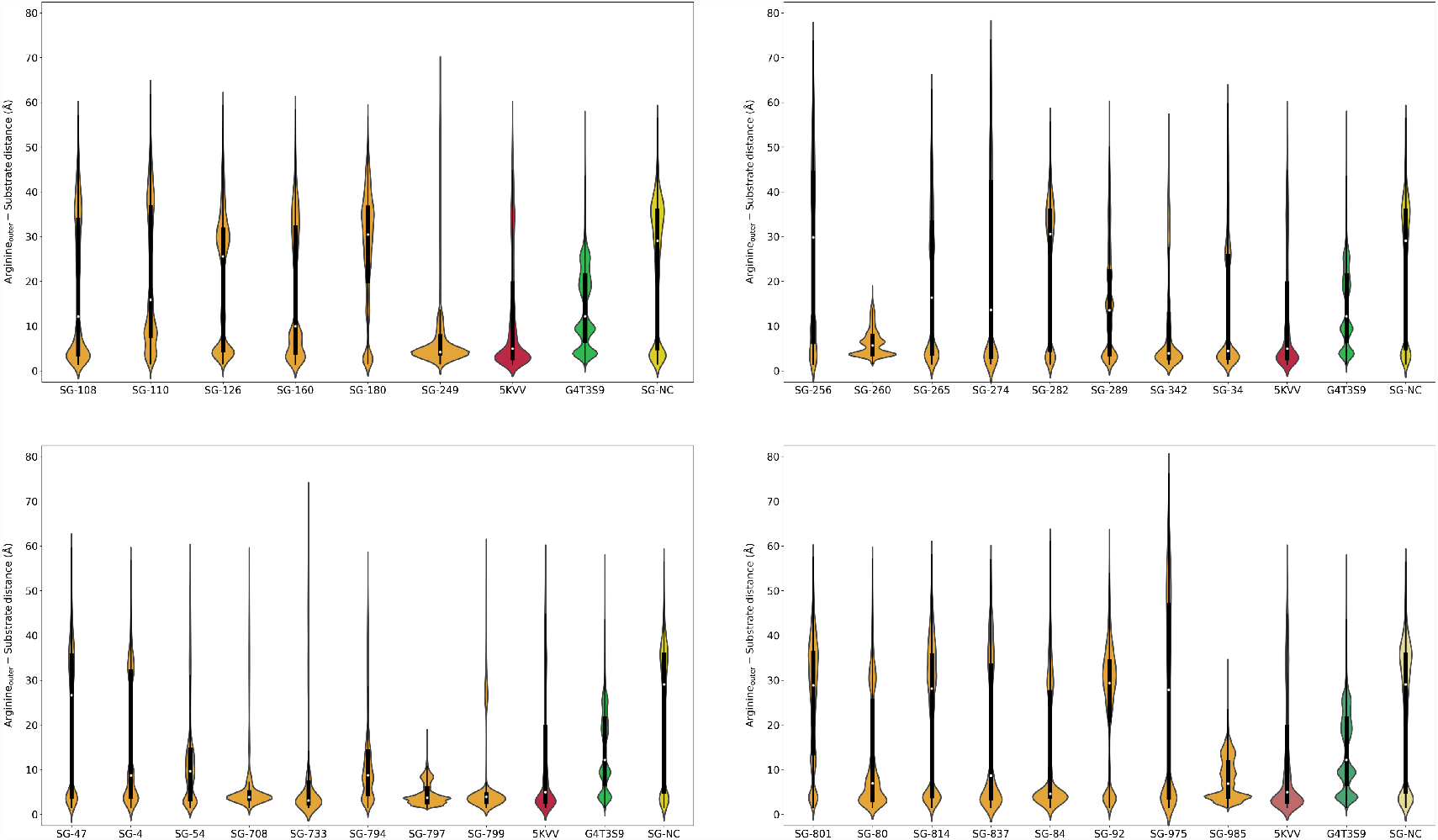
Distribution of the outer Arginine-Substrate 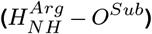 distance of the simulated MDHs. The generated sequences are colored in orange, while the 5KVV control is in red, the G4T3S9 control is in green, and the generated sequence with bad plDDT in the ESMFold prediction is in khaki.

**Supp. Fig. 13.**
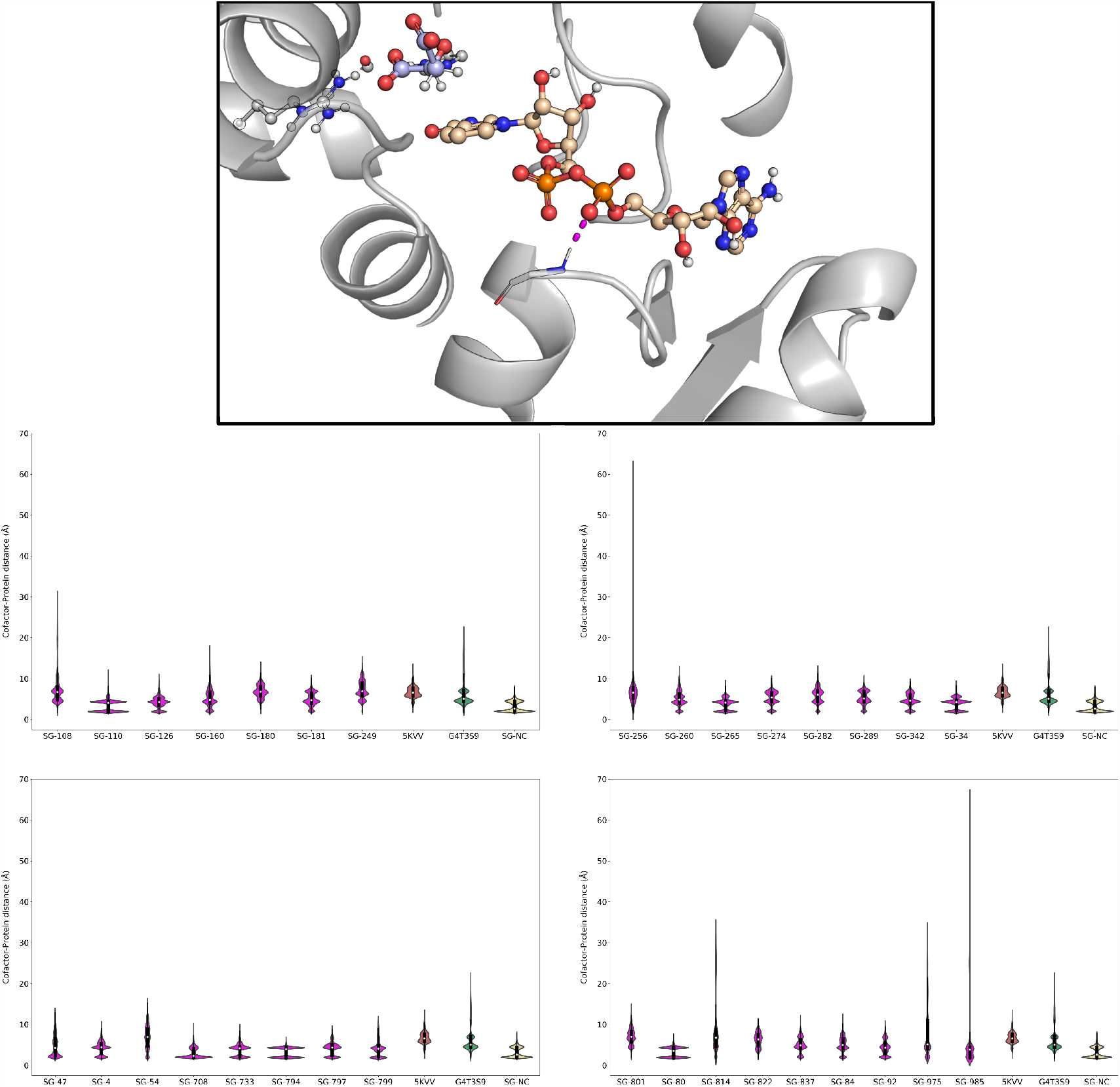
3D representation of the Cofactor-Protein distance in a simulated MDH (top) and distribution of this metric in the simulated MDHs (bottom). The generated sequences are colored in magenta, while the 5KVV control is in red, the G4T3S9 control is in green, and the generated sequence with bad plDDT in the ESMFold prediction is in khaki.

**Supp. Table 2.** Fraction of NAC in the MD simulations. The table contains the percentage of frames from the MD simulation that are NAC. A NAC is considered when the Cofactor-Substrate distance, the Histidine-Substrate distance, the Aspartate-Histidine distance, and the *Arginine*_*inner*_ *− Substrate* distance are ≤ 3.5Å.

| MDH | NAC (replica 1) | NAC (replica 2) | NAC (replica 3) | NAC (replica 4) |
| --- | --- | --- | --- | --- |
| 5KVV | 0.025 | 0.5 | 0.045 | 0.09 |
| G4T3S9 | 0.0 | 0.0 | 0.0 | 0.0 |
| SG-NC | 0.18 | 0.07 | 0.055 | 0.0 |
| SG-108 | 0.705 | 0.305 | 0.11 | 1.655 |
| SG-110 | 0.075 | 0.0 | 0.01 | 0.045 |
| SG-126 | 0.0 | 1.035 | 0.0 | 0.02 |
| SG-160 | 0.005 | 0.375 | 0.165 | 0.005 |
| SG-180 | 0.0 | 0.0 | 0.005 | 0.01 |
| SG-181 | 0.0 | 0.0 | 0.0 | 0.015 |
| SG-249 | 0.0 | 2.725 | 0.75 | 0.02 |
| SG-256 | 0.265 | 1.095 | 0.035 | 0.02 |
| SG-260 | 0.0 | 0.0 | 0.0 | 0.0 |
| SG-265 | 0.08 | 0.36 | 0.47 | 0.185 |
| SG-274 | 0.0 | 0.0 | 0.265 | 0.015 |
| SG-282 | 0.08 | 0.025 | 0.02 | 0.0 |
| SG-289 | 0.055 | 0.12 | 0.935 | 0.28 |
| SG-34 | 0.58 | 1.23 | 0.085 | 0.045 |
| SG-342 | 0.095 | 0.03 | 0.16 | 0.315 |
| SG-4 | 0.015 | 0.025 | 0.025 | 0.02 |
| SG-47 | 0.355 | 0.58 | 0.58 | 0.05 |
| SG-54 | 3.03 | 0.635 | 0.43 | 0.35 |
| SG-708 | 0.005 | 0.0 | 0.005 | 0.055 |
| SG-733 | 0.01 | 0.505 | 2.91 | 1.345 |
| SG-794 | 0.0 | 0.0 | 0.005 | 0.01 |
| SG-797 | 0.02 | 0.045 | 0.05 | 0.335 |
| SG-799 | 0.79 | 0.615 | 0.325 | 0.2 |
| SG-80 | 1.515 | 0.035 | 0.06 | 0.205 |
| SG-801 | 0.0 | 0.01 | 0.0 | 0.0 |
| SG-814 | 0.0 | 0.01 | 0.01 | 0.0 |
| SG-822 | 0.04 | 0.03 | 0.18 | 4.25 |
| SG-837 | 0.91 | 0.11 | 1.68 | 0.32 |
| SG-84 | 0.16 | 0.055 | 0.0 | 0.01 |
| SG-92 | 0.305 | 0.0 | 0.155 | 0.035 |
| SG-975 | 0.1 | 2.6 | 0.0 | 0.485 |
| SG-985 | 0.0 | 0.0 | 0.01 | 0.105 |

